# Mature dentate granule cells show different intrinsic properties depending on the behavioral context of their activation

**DOI:** 10.1101/309906

**Authors:** Angélique Peret, Claire Pléau, Edouard Pearlstein, Thomas Scalfati, Geoffrey Marti, François Michel, Valérie Crépel

## Abstract

The dentate gyrus (DG) plays a crucial role in learning, memory and spatial navigation. Only a small fraction of mature dentate granule cells (mDGCs) is active during behavior, while the large majority remains silent. To date, the properties of this active subset of neurons remain poorly investigated. Using fosGFP transgenic mice, we show *ex vivo* that activated mDGCs, from mice maintained in their home cage, exhibit a marked lower intrinsic excitability compared to the non-activated cells. Remarkably, activated mDGCs, from mice trained in a virtual environment, are more excitable than those from mice maintained in their home cage. Therefore, we show that activated mDGCs display different intrinsic properties and excitable states depending on the context of their activation. We propose that these properties could constitute a neural signature of cell assemblies recruited in different behavioral contexts.

## Introduction

The dentate gyrus (DG), an input region of the hippocampal formation, plays a crucial role in learning, memory and spatial navigation (McNaughton and Morris, 1987; Baker et al., 2016). Memory storage and recall involve a discrete population of cells within the DG region (GoodSmith et al., 2017; Hainmueller and Bartos, 2018; Tonegawa et al., 2015). Only a small fraction of dentate granule cells (DGCs) is active during behavior, while the large majority remains silent (Chawla et al., 2005; Diamantaki et al., 2016; Jung and McNaughton, 1993; Neunuebel and Knierim, 2012; Pilz et al., 2016; Skaggs et al., 1996; Stefanelli et al., 2016; Tonegawa et al., 2015; Hainmueller and Bartos, 2018). Since the discovery of the DG’s ability to generate new neurons throughout life (Aimone et al., 2011), it has been proposed that DG neurogenesis provides a substrate for spatial memory and pattern separation (Aimone et al., 2011; Clelland et al., 2009; Kropff et al., 2015; Leutgeb et al., 2007; McNaughton and Morris, 1987; Nakashiba et al., 2012; Neunuebel and Knierim, 2014; Sahay et al., 2011. Kirschen et al., 2017). This role would be facilitated by the high excitability of newly generated neurons (Aimone et al., 2011; Cameron and Mckay, 2001; Kempermann et al., 1997; Lopez-Rojas and Kreutz, 2016; Ninkovic et al., 2007). On the other hand, recent studies have shown that mature dentate granule cells (mDGCs) play a major role in pattern completion (Nakashiba et al., 2012; Hainmueller and Bartos, 2018) and are required for recall of familiar contexts (Vukovic et al., 2013). However, until now the intrinsic physiological properties of these mature cells remain poorly investigated in contrast to those of adult born DGCs. Notably it is yet unestablished whether mDGCs display uniform intrinsic properties or whether these properties could vary depending on the behavioral context of their activation.

Here, we examined *ex vivo* the intrinsic electrophysiological properties of activated DGCs from mice maintained in their home cage or trained in a virtual reality (VR) environment compared to the non-activated cells. The analysis of intrinsic properties of DGCs activated in these conditions was performed using a fluorescent cellular tagging approach based on a transgenic mouse model in which the synthesis of the fosGFP fusion protein is controlled by the promoter of gene *c-fos*. The latter activity-dependent gene expression is commonly used as a functional readout for the neuronal activation (Barth et al., 2004; Czajkowski et al., 2014) Accordingly, activated neurons during a given behavioral task will express shortly and transiently the fluorescent protein enabling their identification for electrophysiological analysis on acute brain slices (Barth et al., 2004). Our data reveal that activated mDGCs, from mice in their familiar home cage, exhibit different intrinsic properties and a marked lower excitability when compared to the non-activated cells. Remarkably, activated mDGCs from mice trained in virtual environment display a more excitable state when compared to the home cage condition. Therefore, our study shows that mDGCs do not constitute a uniform cell population since they display diverse intrinsic properties depending on the behavioral context of their activation. We propose that these properties could constitute the electrophysiological signature of different cell assemblies involved in specific contexts such as a familiar environment or spatial training.

## Results

### fosGFP^+^ cells display characteristics of mature DGCs in home cage and training conditions

In a first set of experiments, we examined the fraction of DGCs activated in two different behavioral contexts. We studied the percentage of DGCs activated in the home cage condition and during training in a virtual linear track. For the training in a virtual linear track, the task was performed using a virtual reality (VR) system (Dombeck et al., 2010; Hainmueller and Bartos, 2018; Harvey et al., 2009; Ravassard et al., 2013; Schmidt-Hieber and Häusser, 2013) (see Materials and Methods). Head-restrained mice were trained to run back and forth along a 150 cm-long bidirectional virtual linear track that had proximal and distal walls with varying patterns as visual cues for orientation (Figure 1A, B). The mice, which were water restricted throughout the experiment (see Materials and Methods), received a water reward at the end of the track after successfully traversing the full track length. After receiving a reward at the first reward zone, the mice then turned around for running in the opposite direction to receive the next reward at the end of the track. In the first sessions (up to 8), the mice moved slowly and erratically along the track as they learned the task. By 8-10 sessions, the mice moved consistently back and forth along the track, and they received rewards at increasing rates over time (1.14 ± 0.05 reward/min, n = 8 mice), they ran reliably back and forth along the track, and they slowed down before reaching the track end consistent with learning of the task (Figure 1C).

**Figure 1.**
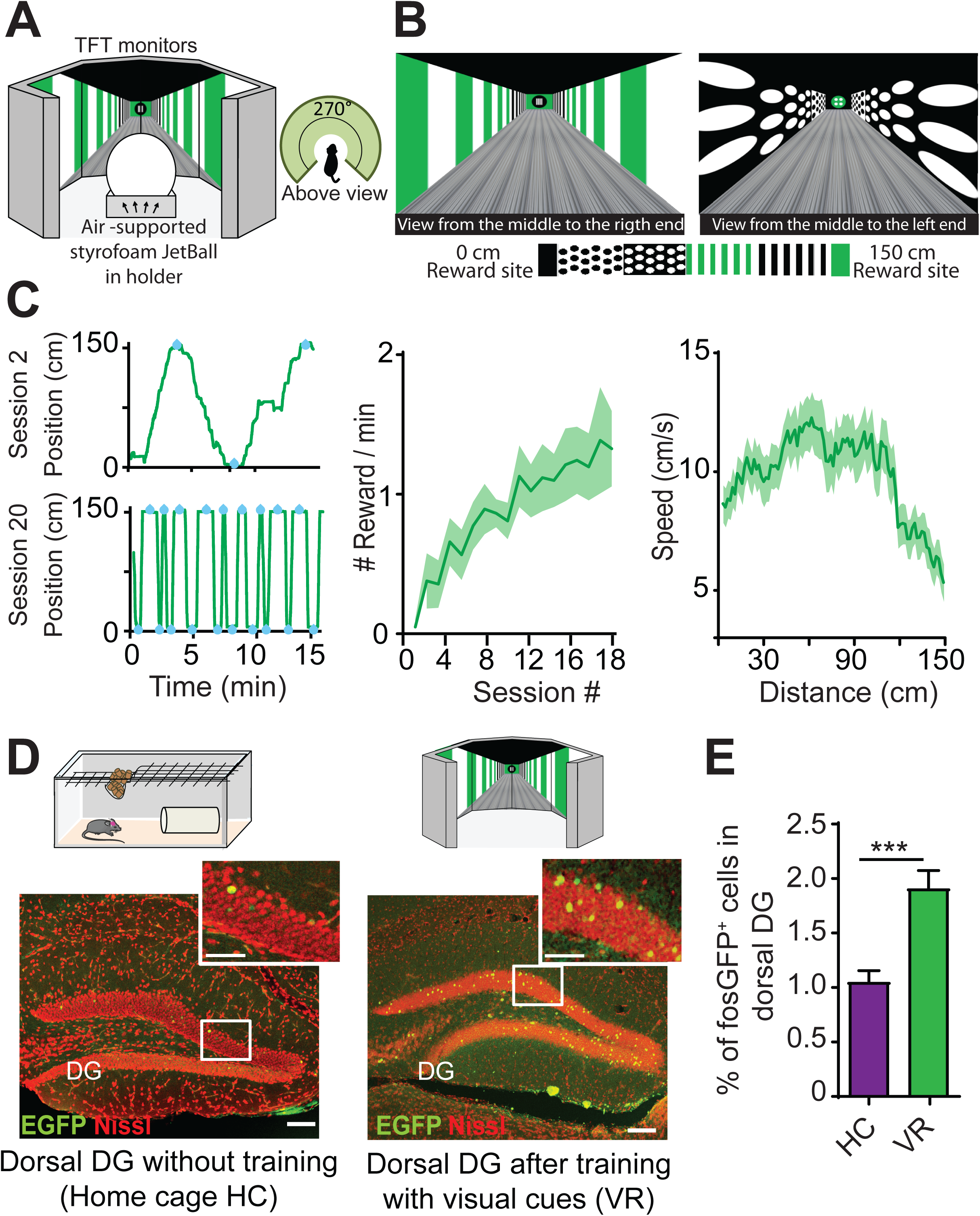
fosGFP^+^ neurons in mice maintained in home cage and trained in VR. (A) Schematic representation of the experimental set-up. Mice were fixed to an arm with a screw through a chronically implanted head-bar. By moving an air-flow-supported styrofoam JetBall, mice were navigating in a 270° virtual environment provided through six TFT monitors surrounding the JetBall (see top view representation of the experimental set-up on top right). Reward was delivered to the animal through a plastic tube. (B) Screenshots from the middle to the right and left ends of the track. The track (150 cm) was divided into four regions with different textures (black dots, white dots, vertical green stripes and vertical black stripes). Water rewards were given at the ends of the track, with available rewards alternating between reward sites. (C) Left: Sample trajectories for an individual mouse on training sessions 2 (top) and 20 (bottom); position is the animal’s location along the 150 cm-long linear track axis and blue dots indicate rewards (left panel). Middle: Rate of rewards vs. session numbers. Right: Running speed of mice as a function of the position on the linear track. (D) On the top, schematic of the different experimental conditions: home cage and training in VR. On the bottom: Nissl staining (red) of dorsal DG with an immunostaining using anti-EGFP antibody (green); scale bar 100 µm. Insets show parts of dorsal DG at higher magnification; scale bar 50 µm. (E) Bar graphs displaying the percentage of fosGFP^+^ cells activated in dorsal DG in mice in home cage (HC) condition and mice trained in VR (VR). In this and following Figures, data are shown as mean ± SEM. *p<0.05, ***p<0.0001, ns, p>0.05.

To analyse the fraction of DGCs activated in the different behavioral contexts (see above), we used a strain of transgenic mice in which the synthesis of fosGFP fusion protein is controlled by the promoter of the activity-dependent immediate early gene (IEG) *c-fos* (see Materials and Methods). These mice enabled an *ex vivo* characterization of activated cells with a recent history of elevated activity in vivo (Barth et al., 2004; Yassin et al., 2010). In VR training conditions, examination of cells expressing fosGFP was performed *ex vivo* in hippocampal slices obtained shortly after the last training session (about 18-20 sessions, see Materials and Methods). Quantification of cells expressing fosGFP was performed in the dorsal hippocampus, which is known to play a crucial role in spatial memory (Moser et al., 1993). When mice were maintained in their home cage, a discreet fraction of DGCs (Prox1-positive, Figure 2C) was activated, since they expressed fosGFP (fosGFP^+^ cells) (1.06 ± 0.1% of DGCs, n = 25 slices, n = 6 mice) (see Figure 1D,E, Materials and Methods). In line with previous observations (Liu et al., 2012; Kirschen et al., 2017; Shevtsova et al., 2017; Stefanelli et al., 2016), we observed a significant higher fraction of fosGFP^+^ cells in the dorsal hippocampus in mice trained in VR (1.92 ± 0.16% of DGCs, n = 19 slices, n = 5 mice, p< 0.0001) (Figure 1D, E).

**Figure 2.**
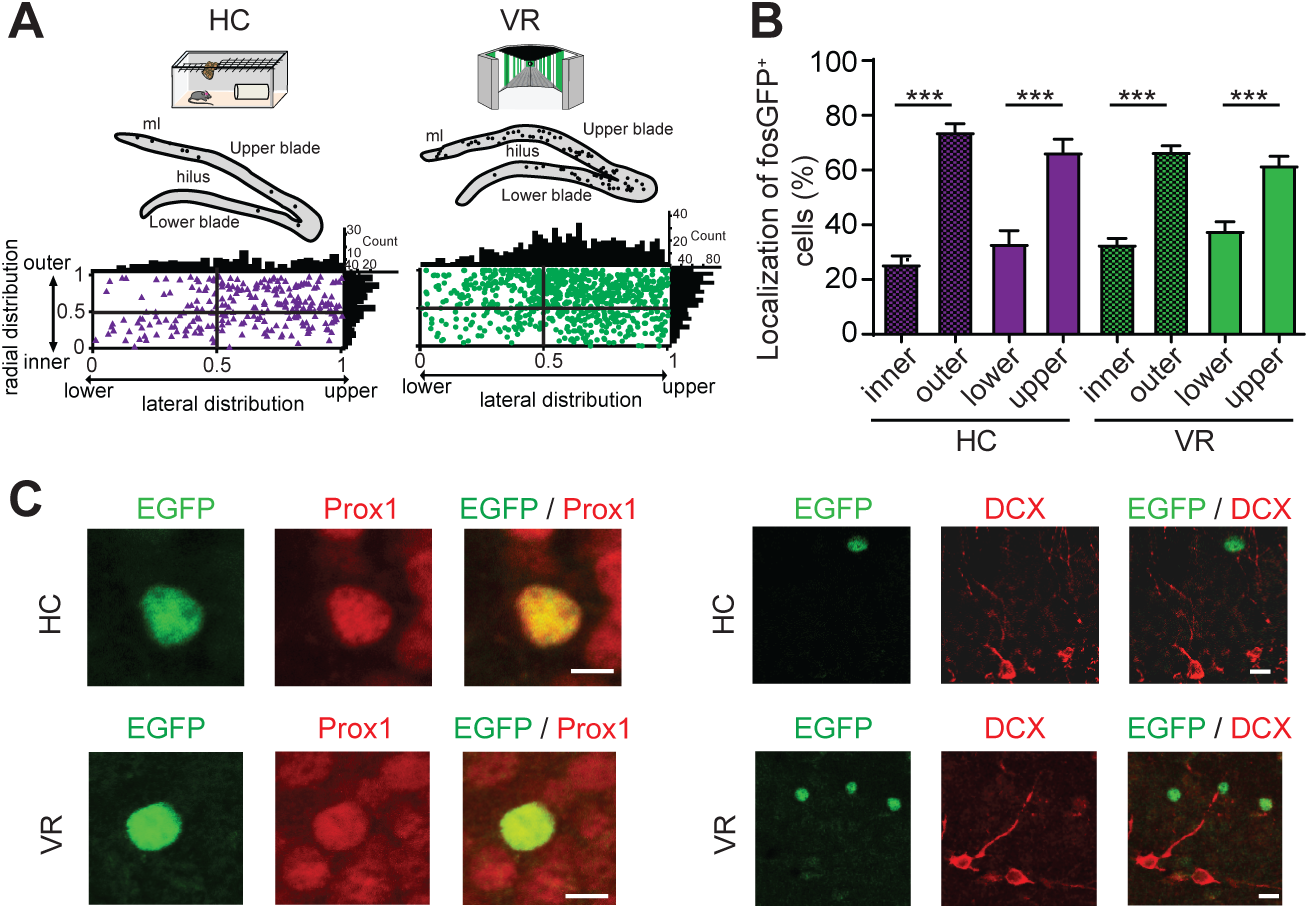
fosGFP^+^ cells display characteristics of mature DGCs. (A) Top: Schematic illustration of DG from confocal image of Figure 1D with fosGFP^+^ cells represented as solid circles. Bottom: Plot diagram representing lateral and radial distributions of fosGFP^+^ from dorsal DG in mice in home cage (HC, triangles) and trained in VR (VR, circles); this cell layer extends laterally from the upper (suprapyramidal) blade to the lower (infrapyramidal) blade and from the outer layer (near the molecular layer) to the inner layer (near the hilus) in a radial direction. The tip of lower blade of granule cell layer is defined as 0 and that of upper blade as 1 in the lateral direction. The border between granule cell layer and the hilus is defined as 0 (inner) and that between granule cell layer and molecular layer (ml) as 1 (outer) in the radial direction. The histograms above and on the right of the square diagram indicate the number of fosGFP^+^ cells in the lateral and radial direction, respectively. (B) Bar graph of the percentage of localization of fosGFP^+^ cells for radial (inner vs outer) and lateral (lower vs upper) distributions in HC and VR conditions. Data are shown as mean ± SEM. ***p<0.0001. (C) Representative double immunostaining of a DGC from HC (top) and from VR (bottom) mice using anti-EGFP antibody associated with anti-Prox1 (left) or anti-DCX antibody (right); scale bar 5 µm.

We then explored the distribution of fosGFP^+^ cells across the DG in dorsal hippocampus in the different experimental conditions. To address this question, we plotted the localization of fosGFP^+^ cells in DG and investigated their lateral and radial distributions. DG cell layer extends laterally from the upper (suprapyramidal) blade to the lower (infrapyramidal) blade and from the outer layer (near the molecular layer) to the inner layer (near the hilus) in a radial direction (Altman and Bayer, 1990). In order to reliably quantify the position across data we used normalized coordinates (see Materials and Methods). We plotted the distribution and the count of fosGFP^+^ cells (Figure 2A, B). Our data showed that for both experimental conditions, a majority of fosGFP^+^ cells were located in the upper blade of dentate granule cell layer (67 ± 4.7%, n = 25 slices, n = 6 mice in home cage condition; 62 ± 3.2%, n = 19 slices, n = 5 mice trained in VR) and near the molecular layer (74 ± 2.9%, n = 25 slices, n = 6 mice in home cage condition; 67 ± 2%, n = 19 slices, n = 5 mice trained in VR) (Figure 2A, B). Furthermore, immunohistochemistry performed with Prox1 antibody confirmed that most fosGFP^+^ cells corresponded to DGCs; we found that more than 95% of these neurons were co-labelled with EGFP and Prox1 antibodies (98.5%, n = 130 cells, n = 5 mice trained in VR; 97.2%, n = 36 cells, n = 3 mice in home cage condition) (Figure 2C). This preferential location near the molecular layer suggested that these fosGFP^+^ DGCs corresponded to mDGCs, since adult-born neurons are preferentially located near the hilus (Zhao et al., 2008). To test that hypothesis, we identify adult-born DGCs and we used an anti doublecortin (DCX) antibody, a specific marker of newly born neurons (Zhao et al., 2008). We did not find any fosGFP^+^ DGC co-labelled with DCX antibodies in the two behavioral contexts (Figure 2C), confirming that fosGFP^+^ cells did not correspond to new-born neurons in our experimental conditions.

Overall, our data reveal that fosGFP^+^ DGCs activated in home cage condition or during training correspond to mDGCs. Furthermore, we observe that training in VR induces an increase in the number of fosGFP^+^ DGCs compared to the home cage condition.

### fosGFP^+^ DGCs display a low intrinsic excitability in mice maintained in home cage

We then asked if intrinsic properties of activated and non-activated DGCs in home cage condition were similar. To address this question, we examined the intrinsic excitability of fosGFP^+^ and fosGFP^−^ DGCs from mice in their home cage was examined *ex vivo* in slices using patch-clamp recordings (see Materials and Methods). We observed a marked lower excitability in fosGFP^+^ DGCs compared with the fosGFP^−^ cells as shown by the f/I plots (see Materials and Methods) (p<0.0001) (Figure 3A). This difference in firing frequency was not associated with a modification of the action potential (AP) threshold, amplitude and half-width (Tables 1, 2). We then analysed the passive membrane properties of fosGFP^−^ (n = 27 cells) and fosGFP^+^ DGCs (n = 17 cells). We observed a lower input resistance (Rin) in fosGFP^+^ DGCs when compared to fosGFP^−^ DGCs (185.36 ± 11.54 MΩ and 336.36 ± 12.74 MΩ in fosGFP+ and fosGFP-DGCs, respectively, p< 0.0001), associated with a higher rheobase (70.59 ± 8.07 pA and 34.81 ± 2.29 pA in fosGFP^+^ and fosGFP^−^ DGCs, respectively, p< 0.0001) (Figure 3B, C) (Tables 1, 2), without a significant change in resting membrane potential (RMP), threshold, amplitude and half-width of AP (Tables 1, 2).

**Figure 3.**
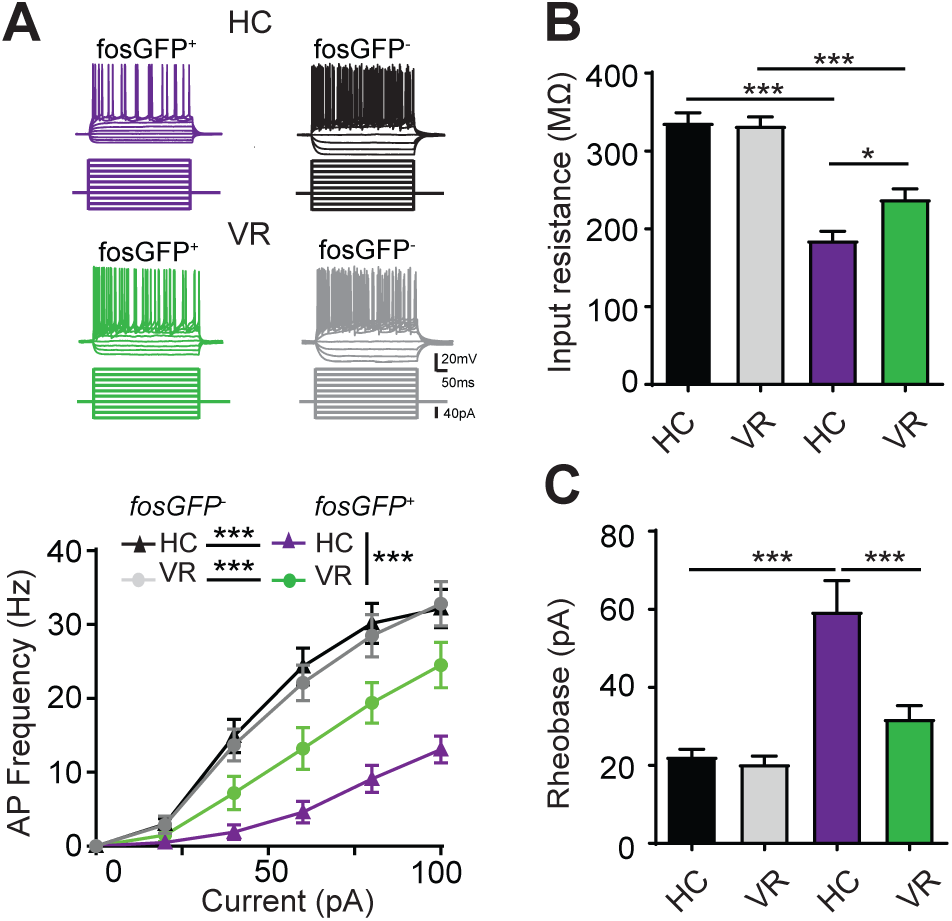
fosGFP^+^ DGCs display different intrinsic properties depending on the behavioral context of their activation. (A) Representative membrane potential variation and action potential discharges in fosGFP^−^ or fosGFP^+^ DGCs in mice in home cage (HC, top) and trained in VR (VR, bottom) evoked by 500 ms current steps varying from −60 pA to +100 pA by 20 pA step increments, at a holding potential of −60 mV. The graph at the bottom represents the discharge frequency/injected current relationship for fosGFP^−^ and fosGFP^+^ DGCs in HC (triangle) and VR (dots) conditions. (B) Bar graphs displaying the input resistance, and (C) rheobase of fosGFP^−^ and fosGFP^+^ DGCs in each condition. Note that the f/I plots, the input resistance, and the rheobase are significantly different when comparing fosGFP^−^ with fosGFP^+^ DGCs, and when comparing fosGFP^+^ DGCs in HC with fosGFP^+^ DGCs in VR.

**Table 1.** Intrinsic electrophysiological properties of fosGFP^−^ and fosGFP^+^ DGCs.

| HC |  |  |  |  |  |  |  |
| --- | --- | --- | --- | --- | --- | --- | --- |
|  |  | R input<br>(MΩ) | Rheobase<br>(pA) | RMP<br>(mV) | AP<br>amplitude<br>(mV) | AP<br>Threshold<br>(mV) | APHW<br>(ms) |
| fosGFP<sup>-</sup> | Mean | 336.40 | 34.81 | -78.30 | 77.65 | -42.26 | 1.22 |
|  | sem | 12.74 | 2.29 | 1.10 | 2.02 | 1.14 | 0.04 |
|  | n | 27 | 27 | 23 | 27 | 27 | 27 |
| fosGFP<sup>+</sup> | Mean | 185.36 | 70.59 | -75.32 | 75.56 | -39.43 | 1.25 |
|  | sem | 11.54 | 8.07 | 1.23 | 2.35 | 1.95 | 0.03 |
|  | n | 17 | 17 | 13 | 17 | 17 | 17 |
| VR |  |  |  |  |  |  |  |
|  |  | R input<br>(MΩ) | Rheobase<br>(pA) | RMP<br>(mV) | AP<br>amplitude<br>(mV) | AP<br>Threshold<br>(mV) | APHW<br>(ms) |
| fosGFP<sup>-</sup> | Mean | 332.40 | 30.83 | -79.25 | 78.39 | -41.39 | 1.18 |
|  | SEM | 11.29 | 2.40 | 1.35 | 1.52 | 0.93 | 0.03 |
|  | n | 24 | 24 | 23 | 24 | 24 | 24 |
| fosGFP<sup>+</sup> | Mean | 237.93 | 43.08 | -77.86 | 73.66 | -40.19 | 1.12 |
|  | sem | 13.48 | 4.99 | 1.83 | 2.84 | 1.78 | 0.06 |
|  | n | 13 | 13 | 11 | 13 | 13 | 13 |

**Table 2.** Intrinsic electrophysiological properties of fosGFP^−^ and fosGFP^+^ DGCs - statistical comparisons.

| Two-way ANOVA p value | R input | Rheobase | RMP | AP amplitude | AP Threshold | APHW |
| --- | --- | --- | --- | --- | --- | --- |
| HC fosGFP<sup>-</sup> / HC fosGFP<sup>+</sup> | <0.0001 | <0.0001 | 0.0795 | 0.6946 | 0.1775 | 0.954 |
| VR fosGFP<sup>-</sup> / VR fosGFP<sup>+</sup> | <0.0001 | <0.0001 | 0.6133 | 0.1411 | 0.2375 | 0.6023 |
| HC fosGFP<sup>-</sup> / VR fosGFP<sup>-</sup> | 0.9006 | 0.1069 | 0.9551 | 0.7459 | 0.9340 | 0.3336 |
| HC fosGFP<sup>+</sup> /<br>VR fosGFP<sup>+</sup> | 0.0174 | 0.0338 | 0.2666 | 0.1274 | 0.7644 | 0.369 |

In conclusion, our data show that fosGFP^+^ DGCs activated in the home cage condition display a lower intrinsic excitability compared to fosGFP^−^ DGCs.

### fosGFP^+^ DGCs display a different excitable state when mice are trained in VR

Our histological analysis has revealed that training in VR induced an important increase in the number of fosGFP^+^ DGCs (see above). In order to study the intrinsic properties of this subset of activated neurons, we examined their firing pattern and intrinsic properties *ex vivo* in slices (see Materials and Methods). In these cells, we observed a higher excitability (Figure 3A) when compared to fosGFP^+^ cells in home cage condition (Figure 3A); this was associated with a higher input resistance (Rin) (237.93 ± 13.48 MΩ in 13 fosGFP^+^ DGCs, p=0.0174) and a lower rheobase (43.08 ± 4.99 pA in 13 fosGFP^+^ DGCs, p=0.0338) (Figure 3B, C), without a significant change in resting membrane potential (RMP), threshold, amplitude and half-width of AP (Tables 1, 2). Remarkably, these fosGFP^+^ DGCs still remained less excitable than fosGFP^−^ cells (Figure 3A, B, C) (Tables 1, 2). This revealed that fosGFP^+^ DGCs trained in VR displayed an “intermediate” state of excitability between DGCs activated in home cage and non-activated DGCs. Finally, our data unveiled that fosGFP^−^ DGCs constituted a homogenous population since they displayed similar excitability and intrinsic properties across experimental conditions (Figure 3) (Tables 1, 2) showing that the diversity of intrinsic properties is restricted to fosGFP^+^ DGCs.

Taken together, our data show that activated DGCs from mice trained in VR display a more excitable state when compared to those from mice in home cage condition.

### fosGFP^+^ DGCs display a different AP waveform and AIS length compared with fosGFP^−^ DGCs

Besides passive and voltage-dependent conductances, intrinsic excitability is tightly regulated by the site of spike initiation in the axon initial segment (AIS) (Araki and Otani, 1955; Bean, 2007; Bender and Trussell, 2012; Coombs et al., 1957; Debanne et al., 2011; Grubb and Burrone, 2010a; Kole and Stuart, 2012; Petersen et al., 2016; Scott et al., 2014; Wefelmeyer et al., 2016). When it is recorded at the somatic level, AP waveform generally comprises two distinct components (Colbert and Johnston, 1996; Coombs et al., 1957; Fuortes et al., 1957; Grace and Bunney, 1983; Häusser et al., 1995), the first peak reflecting the spike initiation in AIS and the second peak the somato-dendritic spike (Bean, 2007). To test a putative modification of AP initiation in AIS in our experimental conditions, we examined the AP waveform in fosGFP^+^ and fosGFP^−^ DGCs using phase plot analysis consisting in the second-derivative of the somatic membrane potential (d^2^V/dt^2^) as a function of somatic membrane potential (Figure 4A) (see Materials and Methods); the second-derivative analysis allowed a good resolution of the two components of the AP phase plot (AIS peak vs. somato-dendritic peak) (Kress et al., 2008; Meeks and Mennerick, 2007). We then evaluated the fraction of fosGFP^−^ and fosGFP^+^ DGCs exhibiting an AP phase plot with one or two peaks (Figure 4A); the first peak was discriminated from the second one using the analysis of the stationary inflection point (Kress et al., 2008). We showed that a majority of fosGFP^−^ DGCs displayed a phase plot with two peaks across all experimental conditions (86%, 25 out of 29 cells in home cage condition; 88%, 23 out of 26 cells in mice trained in VR) (Figure 4B). On the contrary, only around half of fosGFP^+^ displayed a phase plot with two peaks; a significant fraction (p=0.0002) of fosGFP^+^ DGCs displayed a phase plot with a different dynamic, since the first peak could not be detected in this neuronal subset across all experimental conditions (43%, 9 out of 21 cells in home cage condition; 44%, 7 out of 16 cells in mice trained in VR) (Figure 4B). These results reveal a modified axonal component in the AP waveform in fosGFP^+^ cells which could result from a structural modification of the AIS. In order to test this hypothesis, we performed an immunofluorescent labelling for the scaffolding molecule ankyrin-G (AnkG) to quantify AIS length in fosGFP^−^ and fosGFP^+^ DCGs (Figure 5A). Experiments were performed *ex vivo* from mice maintained in home cage condition and from mice trained in VR (see above and Materials and Methods). The analysis was done in DGCs located in the outer part of the molecular layer of the DG (Figure supplement 2B), since we showed that fosGFP^+^ DGCs were preferentially distributed in this area across both experimental conditions (see above and Figure 2). We observed that fosGFP^+^ DGCs displayed a similar shorter AIS length across experimental conditions compared to fosGFP^−^ DGCs (Figure 5C) (23 ± 0.67 µm in fosGFP^−^ n = 47 vs 18.2 ± 0.6 µm in fosGFP^+^ n = 47 in home cage condition, p<0.0001; 24.1 ± 0.63 µm in fosGFP^−^ n = 57 vs 18.9 ± 0.62 µm in fosGFP^+^ n = 57 in mice trained in VR, p<0.0001). This was due to a leftward shift of AIS length distribution towards smaller values (Figure 5B). Interestingly, the shorter AIS length is not associated with a significant relocation of AIS (Figure 5C).

**Figure 4.**
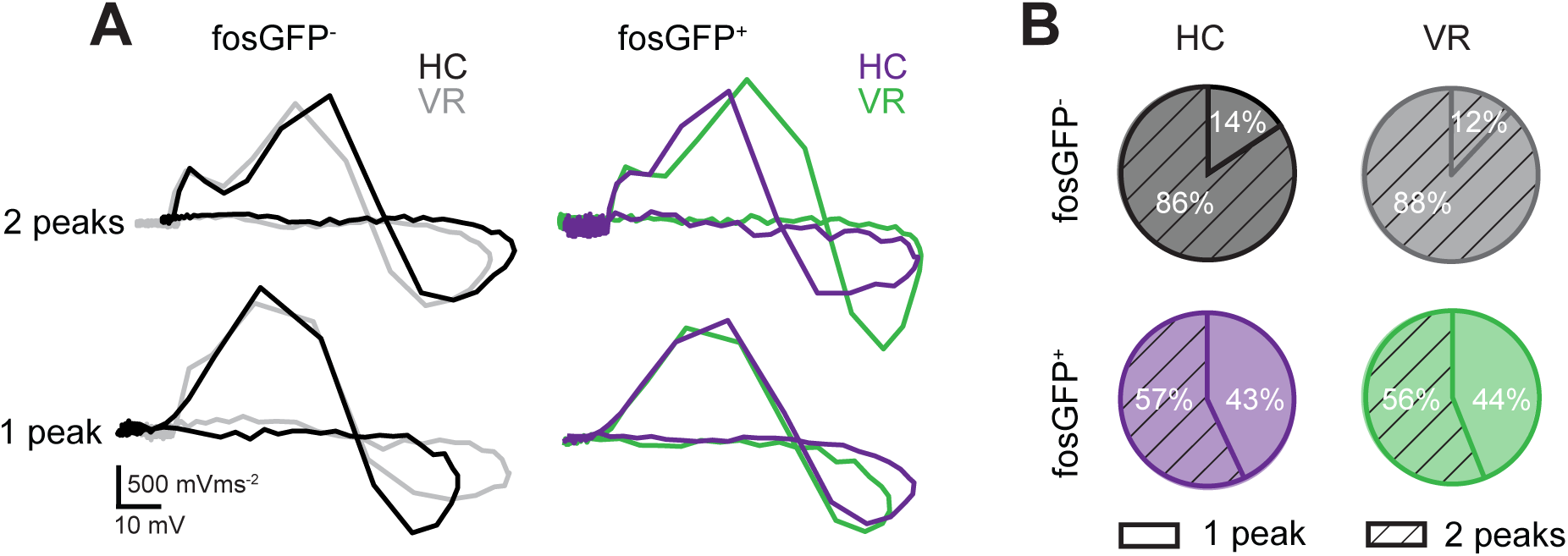
fosGFP^+^ DGCs display a modified AP waveform. (A) Representative superimposed phase plots of the 2nd derivative of the somatic membrane potential (d^2^V/dt^2^) vs. membrane potential (Vm) of action potentials recorded in fosGFP^−^ and fosGFP^+^ DGCs in mice in home cage (HC) and trained in VR (VR); threshold has been taken as reference. Top: phase plots of action potential with two peaks (the first one reflecting the spike initiation in AIS and the second one the somato-dendritic spike). Bottom: phase plots of action potential showing only one peak, the somato-dendritic component. (B) Pie charts of distribution of phase plots with 1 (plain area) or 2 (striped area) peaks in fosGFP^−^ and fosGFP^+^ DGCs in HC and VR conditions. Note that about 90% of fosGFP^−^ DGCs display two peaks, whereas about 50% of fosGFP^+^ DGCs display only one peak.

**Figure 5.**
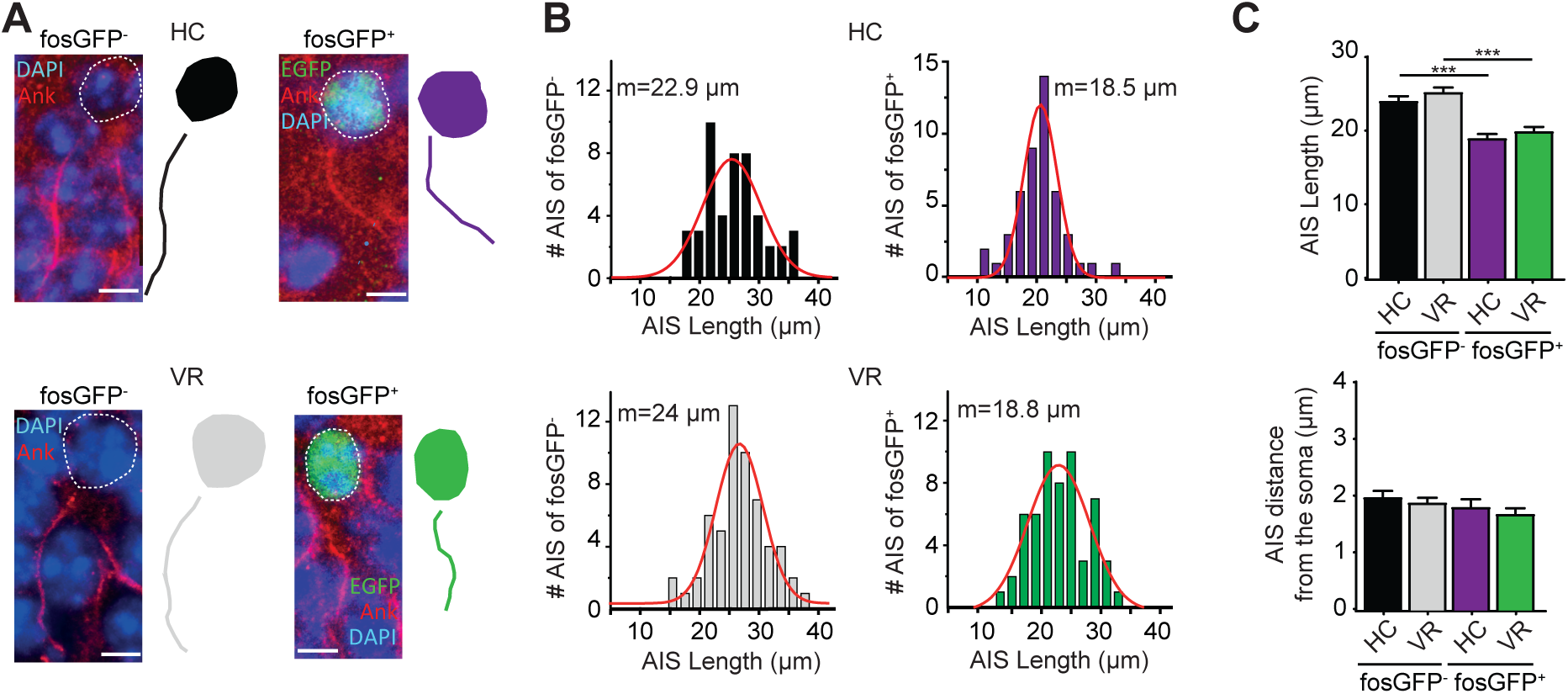
fosGFP^+^ DGCs display a short AIS length. (A) Projection of confocal images of fosGFP^−^ (left) and fosGFP^+^ (right) DGCs showing immunostaining using anti-EGFP, anti-Ankyrin-G antibodies and DAPI, and their Neurolucida reconstructions from mice in home cage (HC) and trained in VR (VR); scale bar, 5 µm. (B) AIS length distribution of fosGFP^−^ (left) or fosGFP^+^ (right) DGCs in HC mice (top) and mice trained in VR (VR, bottom); m indicates the value of the median of the Gaussian fit. Note the shift of AIS length distribution towards smaller values when comparing fosGFP^+^ to fosGFP^−^ cells. (C) Bar graphs displaying the AIS length (left) and the distance from the soma (right) of fosGFP^−^ and fosGFP^+^ DGCs in HC and VR conditions. Note that the length of AIS is significantly shorter in fosGFP^+^ DGCs.

Overall, our data reveal that a significant fraction of fosGFP^+^ DGCs (when compared to fosGFP^−^ DGCs) display a change in the dynamics of their AP waveform associated with a shorter AIS length, which is independent from the behavioral context of their activation.

## Discussion

Our study performed *ex vivo* identifies subsets of mDGCs activated in two different behavioral contexts (home cage condition and training in VR), using a fluorescent cellular tagging approach based on a transgenic mouse model in which the expression of EGFP is controlled by the promoter of the activity-dependent IEG *c-fos* (Barth et al., 2004). Our main finding is that activated mDGCs do not constitute a homogenous neuronal population in terms of intrinsic properties since they display different states of excitability depending on their context of activation. Remarkably, our data also reveal that activated mDGCs always exhibit lower excitability than the non-activated mDGCs across all experimental conditions. In keeping with this, our study shows that AIS length distribution is shifted towards lower values in activated mDGCs. This cellular mechanism could play a role in dampening their basal excitability as previously reported by *in vitro* experiments (Grubb and Burrone, 2010b).

It is well established that the expression of IEGs such as *c-fos* is selectively upregulated in subsets of neurons in specific brain regions associated with learning and memory formation (Minatohara et al., 2015). Notably, many studies have reported that, in these regions, neurons expressing an upregulation of *c-fos* encode and store information required for memory recall (Minatohara et al., 2015; Tonegawa et al., 2015). Therefore, IEGs have been largely used to tag neurons involved in memory function and behavioral tasks. This enables targeting of sparse engram neurons (Tonegawa et al., 2015; Ryan et al., 2015;) to describe their functional features (Barth and Poulet, 2012; Pignatelli et al., 2019). Using fosGFP mice, we identified *ex vivo* a sparse population of DGCs activated in the dorsal hippocampus activated in different behavioral contexts. Notably our findings revealed that training in VR induces a significant increase in the number activated DGCs compared with the home cage condition (Liu et al., 2012; Kirschen et al., 2017; Shevtsova et al., 2017; Stefanelli et al., 2016). These data are in keeping with previous observations showing a sparse recruitment of DGCs during a spatial behavioral experience (Chawla et al., 2005). Remarkably, we showed that, in our experimental conditions, activated DGCs display features of mDGCs since (i) the examination of their radial distribution across the cell layer revealed a majority of fosGFP^+^ cells located near the molecular layer, (ii) we did not observe cells that were co-labelled with EGFP and DCX antibodies, and (iii) they exhibited a mean value of Rin characteristic of mature DGCs (Overstreet-Wadiche and Westbrook, 2006; Dieni et al., 2016; Save et al., 2019). These observations reinforce the notion that mDGCs are not dormant cells (Alme et al., 2010) but comprise active neurons involved in specific tasks (Nakashiba et al., 2012; Vukovic et al., 2013).

We then questioned whether these subsets of mDGCs activated in the home cage condition or during training could have distinct electrophysiological properties. Indeed, there is a growing body of evidence showing that modulation of intrinsic excitability plays a central role in neuronal plasticity and learning processes (Barth, 2007; Daoudal and Debanne, 2003; Sehgal et al., 2013; Zhang and Linden, 2003). Most of the studies investigating a change in intrinsic excitability induced by learning have reported an increase in the firing rate (Daoudal and Debanne, 2003; Epsztein et al., 2011). Surprisingly, we observed an opposite scheme, as activated mDGCs display an overall lower intrinsic excitability than the non-activated cells. Besides this major observation, our findings reveal that activated DGCs do not constitute a homogenous population in terms of intrinsic properties since they exhibit different excitable states depending on the behavioral context of their activation. Indeed, we observed a significant higher excitability in activated DGCs from the mice trained in VR when compared to the cells from mice in home cage condition. Therefore, when comparing intrinsic properties of active neurons, our observations are in keeping with previous observations reporting an increase in excitability during training or learning processes (Barth, 2007; Daoudal and Debanne, 2003; Pignatelli et al., 2019; Sehgal et al., 2013; Titley et al., 2017; Yiu et al., 2014; Zhang and Linden, 2003). We then observed that the difference in excitability is related to the value of input resistance. Indeed, in home cage condition activated mDGCs exhibit a much lower input resistance associated with a lower excitability than the non-activated cells. Input resistance and excitability become significantly higher in activated mDGCs. Interestingly, in both conditions this difference in input resistance is not associated with a significant change in RMP. It is well established that input resistance is set by a variety of leak and voltage-dependent conductances and can be modulated by several intracellular signalling pathways (Brickley et al., 2001; Lesage, 200; Marder and Goaillard, 2006). Therefore, further experiments should be performed to elucidate the mechanisms involved in the setting of input resistance in mDGCs depending on their context of activation.

Another remarkable feature observed is the different dynamics of the phase plot of AP in activated DGCs compared to non-activated cells. Indeed, our data show that nearly all non-activated DGCs displayed a phase plot with two peaks accordingly to previous observations (Kress et al., 2008) across both experimental conditions. On the contrary, an important fraction (almost half) of activated DGCs display a phase plot where the first peak could not be detected by the analysis of the stationary inflection point (Kress et al., 2008). These data, that suggested a change in the axonal component of the AP waveform, could result from a structural modification of AIS. The AIS is a key element involved in the regulation of neuronal excitability since this cellular compartment is responsible for action potential initiation (Araki and Otani, 1955; Bean, 2007; Bender and Trussell, 2012; Coombs et al., 1957; Debanne et al., 2011; Grubb and Burrone, 2010a; Kole and Stuart, 2012; Petersen et al., 2016; Scott et al., 2014; Wefelmeyer et al., 2016). Moreover, it is now well accepted that AIS is not just a rigid structure, but it continuously adapts to its surrounding environment. These changes occur on different time scales ranging from hours to days. AIS modifications can be long-lasting, bidirectional, and are correlated with changes in intrinsic excitability (Evans et al., 2015; Grubb and Burrone, 2010b; Kuba et al., 2010; Wefelmeyer et al., 2016). A structural plasticity of AIS has been reported after several days in auditory nucleus neurons (Kuba et al., 2010). This plasticity, associated with alterations in excitability, contributes to long-term control of AP firing (Grubb and Burrone, 2010a; Kole and Stuart, 2012; Kuba et al., 2010; Wefelmeyer et al., 2015). The precise cellular mechanisms involved are not well understood but recent studies revealed that AIS plasticity depends on calcineurin signalling pathway (Evans et al., 2015). We now show that activated mDGCs display a shorter AIS length than the non-activated cells. As we observed similar reduced AIS length across all experimental conditions, we propose that this difference constitutes a common cellular signature of activated mDGCs independently of their context of activation, which could lead to an overall dampening of excitability via an homeostatic phenomenon (Wefelmeyer et al., 2016). Further experiments should be performed to clarify this point. We hypothesize that this prominent hypo-excitability could reinforce the sparse activity (Chawla et al., 2005; Diamantaki et al., 2016; Jung and McNaughton, 1993; Neunuebel and Knierim, 2012; Pilz et al., 2016; Skaggs et al., 1996; Stefanelli et al., 2016; Tonegawa et al., 2015) and the function of DGCs as coincidence detectors (Schmidt-Hieber et al., 2007).

Taken together, our findings highlight a remarkable feature of activated mDGCs. We found that these neurons show different intrinsic properties depending on their behavioral contexts of activation, besides an overall dampening of excitability. We propose that these properties could constitute the neural signature of cell assemblies involved in specific contexts such as a familiar environment or spatial training.

## Materials and methods

### Ethics

All experiments were approved by the Institut National de la Santé et de la Recherche Médicale (INSERM) animal care and use committee and authorized by the Ministère de l’Education Nationale, de l’Enseignement Supérieur et de la Recherche, following evaluation by a local ethical committee (agreement number APAFIS#9896-201605301121497v11) in accordance with the European community council directives (2010/63/UE).

### Mice

Adult males (n = 91, 100 ± 3 days old, 25.7 ± 0.3 g weight) fosGFP heterozygous mice (B6.Cg-Tg (Fos/EGFP) 1-3Brth/J, Jackson Laboratory, Bar Harbor, ME; RRID = IMSR_JAX:014135) were used for experiments. These mice were generated by fusing the *c-fos* promoter and the *c-fos* coding region, including exons and introns, to the coding region for EGFP, creating a fosGFP C-terminus fusion protein (Barth et al., 2004). All mice were housed in standard conditions (12 hours light/dark cycles at 22 to 24°C, light off at 7:30 a.m., housed one per cage, and food ad libitum) and water restricted (1 ml a day). Mice were handled before recording sessions to limit stress and experiments were performed during the dark cycle.

### Surgical procedures

After the mice being genotyped, a head-bar was implanted. Before the surgery, fosGFP mice were anaesthetized with xylazine (13 mg/kg) / ketamine (66 mg/kg) in 0.9% saline and placed into a stereotactic frame. The skull was exposed and cleaned. Two screws were driven through small holes drilled in the bones and the head-bar was glued to the skull and fixed with bone cement (Heraeus Kulzer GmbH, Hanau, Germany). After 2-3 days of recovery animals were habituated to handling (1-2 days) and were water restricted (approximately 1 ml water/day). If mice weight dropped below 80% of pre-water restriction weight, they had access to food and water ad libitum and were discarded from the study.

### Virtual reality set-up

In the present study we trained mice in virtual reality in order to create a context different from the home environment. Mice were first habituated to be head-restrained on an air-flow-supported Styrofoam ball and trained for 18-20 sessions (30 min per session, 2 sessions per day) (Phenosys GmbH; Berlin, Germany; Figure 1). Animals were running in a 150 cm-long linear track with visual cues on the side-walls (black dots, white dots, vertical green stripes and vertical black stripes) and distal visual cues on both sides provided through six TFT monitors surrounding the animal (JetBall-TFT, Phenosys GmbH), allowing them to identify their position throughout the linear track. Although mice were allowed to move the JetBall in all directions, only the movements along the long axis of the maze were recorded by the system. Upon arrival to the end of the maze, the ball was immobilized by brakes and mice received a small (approx. 4 μl) water drop through a delivery tube; consecutive rewards at the same end were not available. After having obtained the reward, mice were then able to go back by turning the jet-ball to reach the next reward (Figure 1). When the animals performed sufficient number of session (18-20), they were anaesthetised with xylazine (13 mg/kg) / ketamine (66 mg/kg) and sacrificed. Analyses of the speed of the mice along the linear track and quantification of the number of rewards were performed *a posteriori* using custom-developed software written in MATLAB (MathWorks, Natick, MA).

### Quantification of fosGFP^+^ cells

The examination of cells expressing fosGFP (fosGFP^+^) was performed in hippocampal slices from animals maintained in their home cage, or animals that achieved the last VR training session. Mice were deeply anesthetized with xylazine (13 mg/kg) / ketamine (66 mg/kg) prior to decapitation. The brain was then rapidly removed and hippocampi were dissected and fixed overnight (home cage: n = 6 mice, trained in VR: n = 5 mice). Transverse 80 µm thick slices were cut using a Leica VT1000S vibratome (Leica Microsystems, Wetzlar, Germany). Slices were then permeabilized in blocking solution containing 5% normal goat serum (NGS, Sigma-Aldrich, Merck KGaA, Darmstadt, Germany) in 0.5% Triton for 1 hour at room temperature. Slices were then incubated with the polyclonal rabbit anti-EGFP antibody (Fisher Scientific, Hampton, NH) at 1:1000 in 5% NGS in 0.5% Triton x-100 overnight at 4°C. Slices were then incubated for 2 hours with the Alexa Fluor 488 secondary antibody (Invitrogen, Carlsbad, CA, 1:500) and counterstained with Neurotracer fluorescent Nissl (Invitrogen; RRID:AB_2620170) at 1:250 then coverslipped in fluoro-gel (Electron Microscopy Sciences, Hatfield, PA). Fluorescent images were acquired using a confocal microscope (TCS SP5 X, Leica Microsystems) and image analysis was assessed by using ImageJ (NIH). We confirmed that expression of the fosGFP transgene correlated with the synthesis of the endogenous c-fos protein using a c-fos specific antibody (Synaptic Systems GmbH, Göttingen, Germany; RRID: AB_2106755, see below); we found that almost all fosEGFP^+^ neurons were co-labelled with EGFP and c-fos antibodies (100 %, n = 18 cells, n = 2 mice in home cage condition; 89.7 %, n = 146 cells, n = 3 mice trained in VR) (Figure supplement 1). The density of fosGFP^+^ cells was quantified (as cells.mm^-3^) by calculating DG area and counting the number of GFP^+^ cells in each slice. The percentage of fosGFP+ DGCs was estimated by reporting the density of fosGFP+ neurons to the total density of granule cells (i.e. 967 442 cells/mm3), which was calculated from the total number of granule cells in the DG (Buckmaster and Wen, 2011) and the volume of the DG (van Praag et al., 1999). Spatial distribution of fosGFP^+^ neurons were examined through the DG cell layer. This neuronal layer extends laterally from the upper (suprapyramidal) blade to the lower (infrapyramidal) blade and from the outer layer (near the molecular layer) to the inner layer (near the hilus) in a radial direction (Altman and Bayer, 1990; Muramatsu et al., 2007). In the lateral axis, we defined the tip of DG lower blade as 0 and the tip of DG upper blade as 1. Similarly, we defined in the radial direction the border between granule cell layer and the hilus as 0 and the border between granule cell layer and molecular layer as 1. Therefore, each fosGFP^+^ DGC was assigned two values between 0 and 1 corresponding to its respective position along the radial and lateral axis. The distribution of fosGFP^+^ was then plotted (Figure 2A). The quantification of fosGFP^+^ cells was performed on the dorsal hippocampus, as this region has been shown to be tightly associated with learning and memory (Fanselow and Dong, 2010)

### Immunohistochemistry

For EGFP, cfos and Prox1 staining, slices were fixed then permeabilized in blocking solution containing 5% normal goat serum (NGS) (Sigma-Aldrich) and 0.5% Triton for 1 hour at room temperature. The slices were incubated with polyclonal chicken anti-EGFP (Abcam, Cambridge, UK; RRID:AB_300798) at 1:1000 and either with polyclonal Guinea pig anti-c-fos antibody (Synaptic Systems; RRID: AB_2106755) at 1:500 or polyclonal rabbit anti-Prox1 antibody (Millipore; Merck KGaA, Darmstadt, Germany, RRID:AB_177485) at 1:2000 or polyclonal rabbit anti-DCX antibody (Abcam; RRID:AB_2088478) at 1:1000 in 5% NGS in 0.5% Triton overnight at 4°C. Slices were then incubated for 2 hours with secondary antibodies Alexa488 or Alexa555 (Invitrogen 1:500) and then coverslipped in fluoro-gel.

Specific protocol has been used for Ankyrin G immunostaining (NeuroMab; UC Davis/NIH NeuroMab Facility, Davis, CA; RRID: AB_10673030). Trained mice were first deeply anesthetized with an i.p. injection of xylazine (13 mg/kg) / ketamine (66 mg/kg), then briefly intracardiacally perfused (flowrate: 8–10 ml/min for 2–5 min) with cold 1X phosphate buffer (PBS, pH 7.4), followed by 10 min of cold 1% antigenfix (Diapath, Martinengo, Italy). Brains were removed carefully and post-fixed in antigenfix for 24 h at 4°C and then cryopreserved in 20–30% sucrose / PBS at 4°C. Brains were allowed to completely sink to the bottom of the container before sectioning. Then, they were embedded in OCT compound (Tissue-Tek R, TedPella Inc., Redding, CA) and placed at −80°C. Frozen brains were sectioned transversally into 30 μm thick slices at −20°C to −25°C using a Leica CM 3050S cryostat (Leica Microsystems). Floating brain sections were washed with 1X PBS, and then pre-incubated with permeabilizing agent containing 0.5% Triton, 0.2% Tween-20 and 3% BSA. Then slices were washed with 0.2% Tween-20 and incubated with a blocking buffer consisting of 5% NGS (Sigma-Aldrich) and monoclonal mouse anti-ankyrinG antibody (1:500) (NeuroMab) overnight at 4°C. Slices were washed and incubated with biotin-conjugated secondary antibody (1:500) (Invitrogen) followed by incubation with Cy3-conjugated streptavidin (1:500) (Sigma Aldrich), then coverslipped with mounting medium containing DAPI (Vector laboratories, Burlingame, CA). Image acquisition with a Leica confocal microscope was performed to associate each AIS labelled with anti-ankyrinG antibody with the corresponding fosGFP^+^ cell via EGFP staining and fosGFP^−^ cell via DAPI staining. These images were also used for the reconstruction of the soma and the AIS with Neurolucida software. Image acquisition with a Leica confocal microscope (40 X oil immersion objective; using a laser scanning with appropriate excitation and emission filters) was performed to associate each AIS labelled with anti-ankyrinG antibody with the corresponding fosGFP^+^ cell via EGFP staining and fosGFP^−^ cell via DAPI staining. These images were also used for a 3D-reconstruction of the soma and the AIS with Neurolucida software. The ‘start’ and ‘end’ positions of AIS were the proximal and distal axonal points, respectively, at which the fluorescence profile dipped to 33% of its maximum peak (Figure supplement 2A) (Grubb and Burrone, 2010a).

### Acute slice preparation

Hippocampal slices were prepared from heterozygous fosGFP mice. The slices were cut 45 min after the last training session in VR (18-20 sessions). Animals were deeply anesthetized with xylazine (13 mg/kg) / ketamine (66 mg/kg) prior to decapitation. The brain was then rapidly removed, and transverse 350 µm thick slices were cut using a Leica VT1200S vibratome in ice-cold oxygenated (95% O_2_ and 5% CO_2_) modified artificial cerebrospinal fluid (ACSF) containing the following (in mM): 132 choline, 2.5 KCl, 1.25 NaH_2_PO_4_, 25 NaHCO_3_, 7 MgCl_2_, 0.5 CaCl_2_, and 8 D-glucose. Slices were transferred to rest at room temperature in oxygenated (95% O_2_ and 5% CO_2_) solution of ACSF containing the following (in mM): 126 NaCl, 3.5 KCl, 1.2 NaH_2_PO_4_, 26 NaHCO_3_, 1.3 MgCl_2_, 2.0 CaCl_2_, and 10 D-glucose, pH 7.4.

### Electrophysiological recordings

Individual slices were transferred to an upright microscope and neurons were visualized with infrared differential interference contrast microscopy (SliceScope Pro 3000M, Scientifica, Uckfield, UK). Slices were placed in a submerged chamber and perfused with oxygenated ASCF (30-32°C) at a flow rate of 2 to 3 ml/min. Whole cell recordings of DGCs were obtained using patch clamp technique. Glass electrodes (resistance 6-8 MΩ) were filled with an internal solution containing the following (in mM): 130 KMeSO_4_, 5 KCl, 10 4-(2-hydroxyethyl)-1-piperazi-methanesulfonic acid, 2.5 MgATP, 0.3 NaGTP, 0.2 ethyleneglycoltetraacetic acid, 10 phosphocreatine, and 0.3 to 0.5% biocytin, pH 7.25. Access resistance ranged between 15 and 30 MΩ, and the results were discarded if the access resistance changed by >20%. Whole cell recordings were performed in current-clamp mode using a Multiclamp 700B amplifier (Molecular Devices, Sunnyvale, CA). Data were filtered at 2 kHz, digitized (20 kHz) with a Digidata 1440A (Molecular Devices) to a personal computer, and acquired using Clampex 10.1 software (PClamp, Molecular Devices). The recordings alternated between targeting fosGFP^−^ and fosGFP^+^ neurons during the course of an experiment. To avoid EGFP bleaching, the tissue was illuminated for a short period of time, typically around 5-10 s, to focus and record the image of targeted neuron.

Electrophysiological parameters were tested in whole-cell patch-clamp, when resting membrane potential (RMP) was stable for at least 5 min. Electrophysiological parameters were measured from responses to step current injections (500 ms duration) increasing from negative to positive values, and applied from a fixed membrane potential of −60 mV. Input resistance (Rin) was determined by plotting the membrane potential variations induced by hyperpolarising 500 ms steps of current (from −60 pA to 0 pA) and measuring the slope of the fitted linear regression curve. Firing frequency was studied by injecting 500 ms pulses of depolarizing current (I: from 20 pA up to 100 pA) into the cell and plotting the spike frequency (f) as a function of the current intensity (f/I plot). The AP waveform was analysed on the first AP elicited at suprathreshold potential (Mini Analysis Program, Synaptosoft, Decatur, GA). AP threshold was defined as the membrane potential when dV/dt > 10 mV/ms (Kress et al., 2008; Naundorf et al., 2006). AP amplitude was the difference between the AP threshold and the AP peak. AP width was measured from the half-height of the AP amplitude. Second-derivative (denoted d^2^V/dt^2^) of the somatic membrane potential was calculated using Clampfit analysis tools. Phase plots were constructed by plotting d^2^V/dt^2^ as a function of somatic membrane potential. Stationary inflection point refers to the point where the second derivative displays a slope of zero (Kress et al., 2008).

### Statistical analysis

All values are given as means ± SEM. Statistical analyses were performed using Graphpad Prism 7 (GraphPad Software, La Jolla, CA). The normality of data distribution was assessed using the Shapiro-Wilk normality test. For comparison between two groups with normal distribution, the two-sample unpaired Student’s t test was used, otherwise we used the unpaired Mann–Whitney test. To investigate the relationship between two parameters, the Pearson’s correlation test was used. All tests were two-sided. For the comparison of multiple groups of two factors, we used two-way ANOVA test. Bonferroni post hoc test was used when adequate. When data were not normally distributed, the unpaired Mann–Whitney rank-sum test was used. The χ^2^ or the Fisher’s exact tests were used to compare proportions. Statistics are provided in the text and tables. The level of significance was set at P < 0.05; exact P values are given, unless P < 0.0001 or P > 0.9999. Analyses were performed blind to experimental groups.

## Acknowledgments

We thank B. Poucet, D. Robbe, A. Malvache, J. Epsztein, J. Koenig, I. Bureau, R. Cossart, A. Represa, R. Khazipov, S. X. Ho, and T. Marissal for helpful comments on this manuscript; and S. Varpula, S. Moussa, T. Tressard, and L. Petit for technical assistances. S. Corby-Pellegrino for heading the animal house facility. This work was supported by the Institut National de la Santé et de la Recherche Médicale (INSERM), Aix-Marseille Université (AMU), the Agence Nationale de la Recherche (ANR) (ANR 13-BSV4-0012-02 to V.C.), and the french Ministère de l’Enseignement Supérieur et de la Recherche (MESR to C.P.).

---

**Supplementary figure 1.**
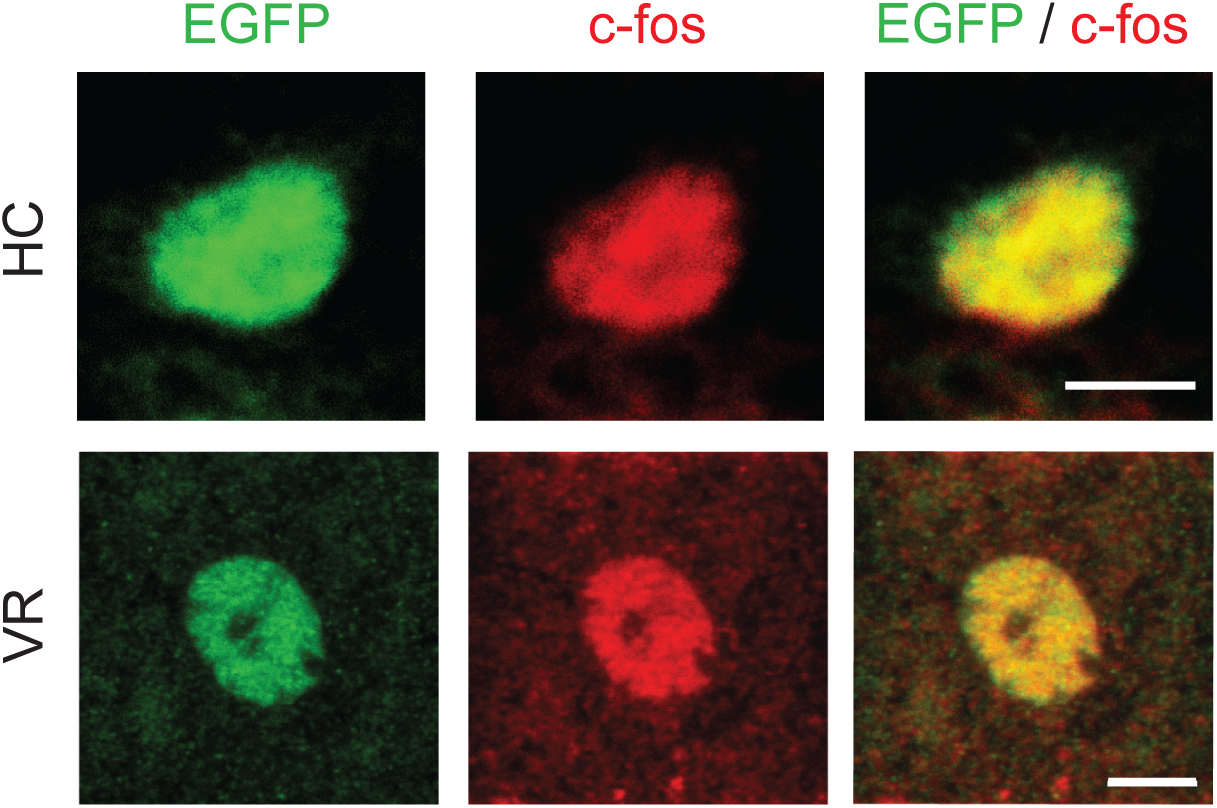
Representative double immunostaining using anti-EGFP antibody associated with anti-c-fos antibody in mice in home cage (HC) and trained in VR; scale bar 5 μm.

**Supplementary figure 2.**
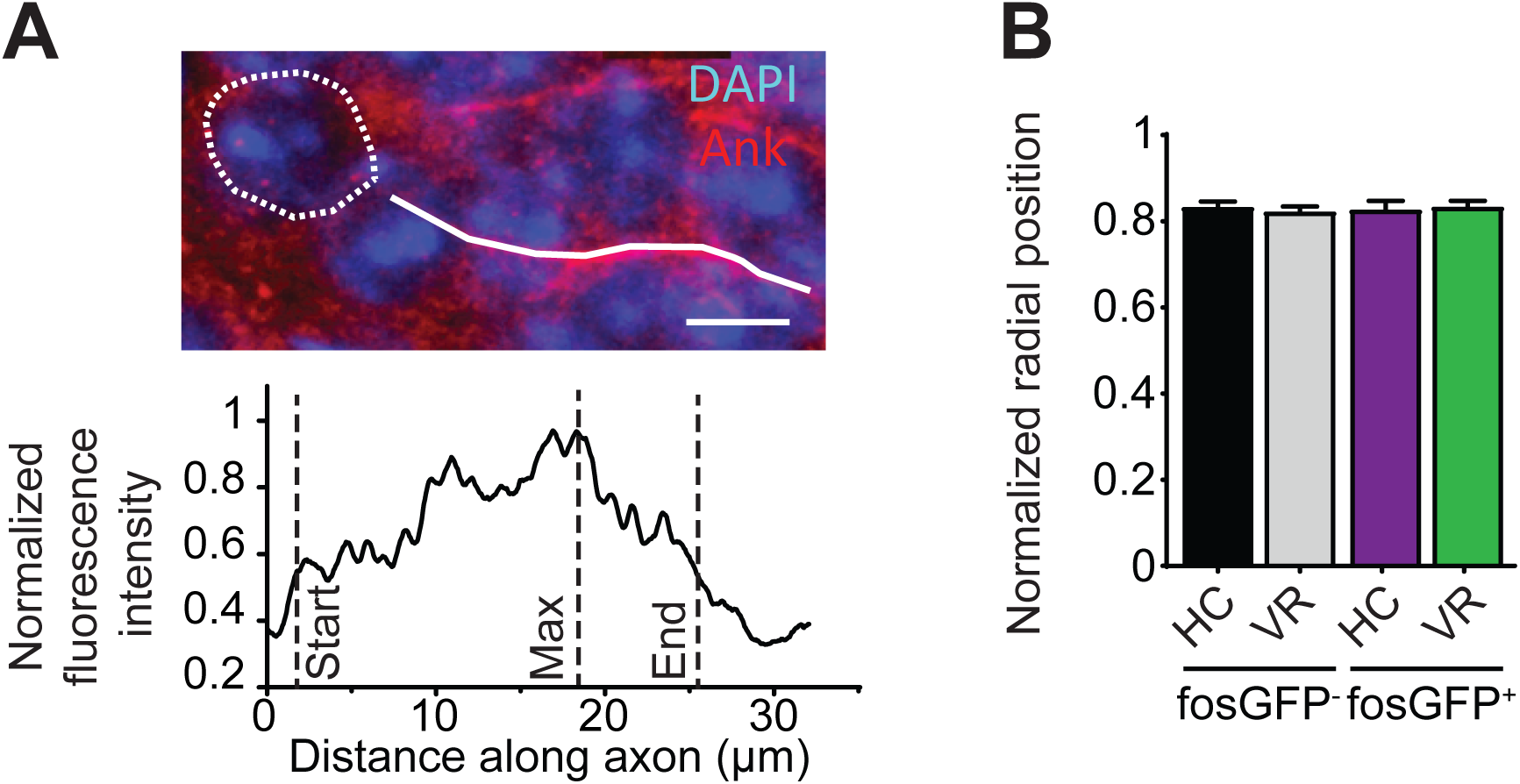
(A) Top, projection of a confocal image showing immunostaining using anti-Ankyrin-G antibodies and DAPI and the superimposed Neurolucida reconstructions of the soma (dotted line) and axon initial segment (solid line); scale bar, 5 µm. Bottom, fluorescence intensity along the axon initial segment. (B) Bar graphs displaying the normalized radial position of fosGFP^−^ and fosGFP^+^ cells used to analyse AIS length and AIS location from the soma in mice in home cage (HC) and trained in VR.

